# BRET-Based Mitochondrial Subcompartment Localization Biosensors

**DOI:** 10.1101/2025.10.27.684728

**Authors:** Abdulrasheed O. Abdulrahman, Mathias Lescuyer, Rami El Zein, Erika Cecon, Julie Dam, Bernard Masri, Ralf Jockers

**Affiliations:** Université Paris Cité, Institut Cochin, INSERM, CNRS, 75014 Paris, France

**Keywords:** Mitochondria, Biosensors, BRET, Bax, VDAC, Drp1

## Abstract

The last decade has witnessed a marked increase in interest in mitochondria, whose dysfunction leads to the development of multiple diseases. Mitochondria are unique as they are highly compartmentalized organelles that are composed of two closely apposed membranes and whose biological function relies on the precise localization of nuclear-encoded proteins in distinct mitochondrial subcompartments. Here we developed a series of bioluminescence resonance energy transfer (BRET)-based localization biosensors to monitor the precise localization of proteins in different mitochondrial subcompartments (outer and inner membrane, intermembrane space at the inner boundary membrane, crista lumen and matrix) with a high spatial resolution (1-10 nm). These biosensors detected the correct localization and orientation of TOM20, TOM22, VDAC1, MICU1, ATP5F1C, OTC, and SIRT2/3 proteins in their respective subcompartments, as well as the translocation of BAX and Drp1 from the cytosol to mitochondria in intact cells. The localization sensors provide non-invasive tools to monitor protein localization and translocation to mitochondria in real-time with nanometer resolution in intact cells submitted to various stressors.

## Introduction

Mitochondria are complex organelles found in virtually all eukaryotes except Monocercomonoides. They are recognized as the primary sites of cellular respiration, where energy stored in nutrients is converted into adenosine triphosphate (ATP) [1]. Apart from their role in energy production, mitochondria are also involved in regulating cellular metabolism, calcium homeostasis, and apoptosis, making mitochondria currently the most studied intracellular compartment [2,3]. Importantly, mitochondrial dysfunction is linked to a number of human pathologies, including neurodegenerative diseases, cancer, and metabolic disorders. Mitochondria are unique as they are composed of two membranes, the Outer Mitochondrial Membrane (OMM), which is in direct contact with the cytosol, and the Inner Mitochondrial Membrane (IMM). The IMM is subdivided by cristae junctions into two structurally, and functionally distinct domains, the Cristae Membrane (CM), containing the respiratory chain generating the proton gradient necessary for ATP production, and the Inner Boundary Membrane (IBM) [4]. The space between OMM and IMM is called inter-membrane space (IMS), the space generated by the CM, cristae lumen (CLU), and the space inside the IMM, matrix (MX). Primary mitochondrial functions, such as ATP production, reactive oxygen species regulation, and metabolite transport, are highly dependent on the precise organization of these sub-compartments [5] [6]. Mislocalization of mitochondrial proteins has been implicated in a spectrum of diseases, including primary hyperoxaluria type 1, neurodegenerative diseases, and cancer [7–9].

A range of techniques have been applied to determine the localization of proteins in mitochondria, such as proteomics techniques coupled to mass spectrometry detection and subcellular fractionation, followed by western blot detection [10]. The main limitations of these techniques are contamination from other compartments, the challenge to resolve mitochondrial subcompartments, the lack of dynamics, and the availability of specific antibodies for the protein(s) of interest.

Microscopic techniques offer more reliable spatial information due to the preservation of the intact cellular environment [11]. Standard fluorescence microscopy is well adapted to determine the localization of proteins in mitochondria, but is unable to determine the localization at the subcompartment level due to the resolution limit of ∼200–250 nm [12]. While high-resolution microscopy techniques such as Stimulated Emission Depletion (STED) and Single-Molecule Localization (e.g., PALM, STORM, SMLM) represent significantly improved spatial resolution (down to ∼20–50 nm) [12], Transmission electron microscopy (TEM) remains currently the technique with the highest spatial resolution (below 0.1 nm), clearly resolving mitochondrial subcompartments[13]. TEM is, however, time-consuming, limited to fixed cells, only semi-quantitative, and the detection of proteins by immunogold labeling depends on the availability of suitable antibodies [14,15].

Alternative techniques relying on molecular proximity, such as Bi-Genomic Mitochondrial-Split-GFP have been applied to the detection of mitochondrial proteins [16]. Energy transfer-based techniques represent another promising approach to probe molecular proximity. They rely on the non-radiative transfer of energy from an energy donor to an energy acceptor when the donor-acceptor pair is in close proximity (1 to 10 nm). Depending on the nature of the energy donor, either a fluorescent protein (or fluorophore) or a luciferase enzyme, we distinguish between Fluorescence or Bioluminescence Resonance Energy Transfer (FRET or BRET). Typical energy donors of BRET assays are luciferase enzymes, such as Renilla luciferase (Rluc) and NanoLuc (Nluc) [17], which emit light in the presence of their respective substrate and oxygen.

Commonly used energy acceptors are GFP variants, e.g. YFP. Energy transfer techniques are widely used to study protein-protein interactions and conformational changes in living cells [22,23]. Molecular proximity of two proteins of interest can be measured by designing a fusion protein between protein 1 and the energy donor and another fusion protein between protein 2 and the energy acceptor. Three examples of mitochondrial protein-protein interactions [24–26] and one of proteins bridging mitochondria and the endoplasmic reticulum (ER) [27] have been reported previously. Alternatively, the localization of proteins in a specific cellular compartment can be monitored by so-called ‘bystander localization sensors’, where a signal peptide for a specific subcellular compartment is fused to the energy donor or acceptor [28]. Such sensors have been designed for the plasma membrane, endosomes, the ER, the Golgi, nucleus, and extracellular vesicles, but not for the different mitochondrial compartments [28–30]. Apart from the highly relevant detection range of 1 to 10 nm to discriminate between mitochondrial subcompartments, energy transfer techniques allow to collect additional information on the orientation of transmembrane-spanning proteins due to the specific fusion of the energy donor/acceptor pair and follow molecular interactions in a dynamic manner in intact cells, an important feature given the highly dynamic nature of mitochondria.

In the present study, we report the development of BRET-based ‘bystander localization sensors’ for the precise detection of mitochondrial proteins in a specific sub-compartment. The localization sensors were extensively characterized and validated through immunofluorescence and luminescence imaging and benchmarked in various BRET assays with mitochondrial marker proteins located in different subcompartments at steady-state and upon apoptosis induction.

## Results and Discussion

### Design of mitochondrial BRET localization sensors

In this study, we developed four localization sensors to probe different mitochondrial sub-compartments, the OMM sensor probing the cytoplasmic surface of the outer mitochondrial membrane (OMM), the inner boundary membrane (IBM) sensor probing the IMS at the level of the IBM, the cristae lumen (CLU) sensor probing the IMS at the level of the cristae and the MX sensor probing the matrix (Figure 1). Localization sensors were designed by fusing BRET donor and acceptor proteins to previously described small sequence stretches of mitochondrial proteins known to target fusion proteins to specific mitochondrial subcompartments (Table 1). The highly sensitive Nluc enzyme was used as energy donor, and the yellow fluorescent protein (YFP) as energy acceptor. Two types of BRET assays were designed, the first, monitoring the energy transfer between the Nluc- and YFP-fused localization sensors, the second, monitoring the energy transfer between Nluc-fused full-length subcellular marker proteins and the YFP-fused localization sensors.

**Figure 1:**
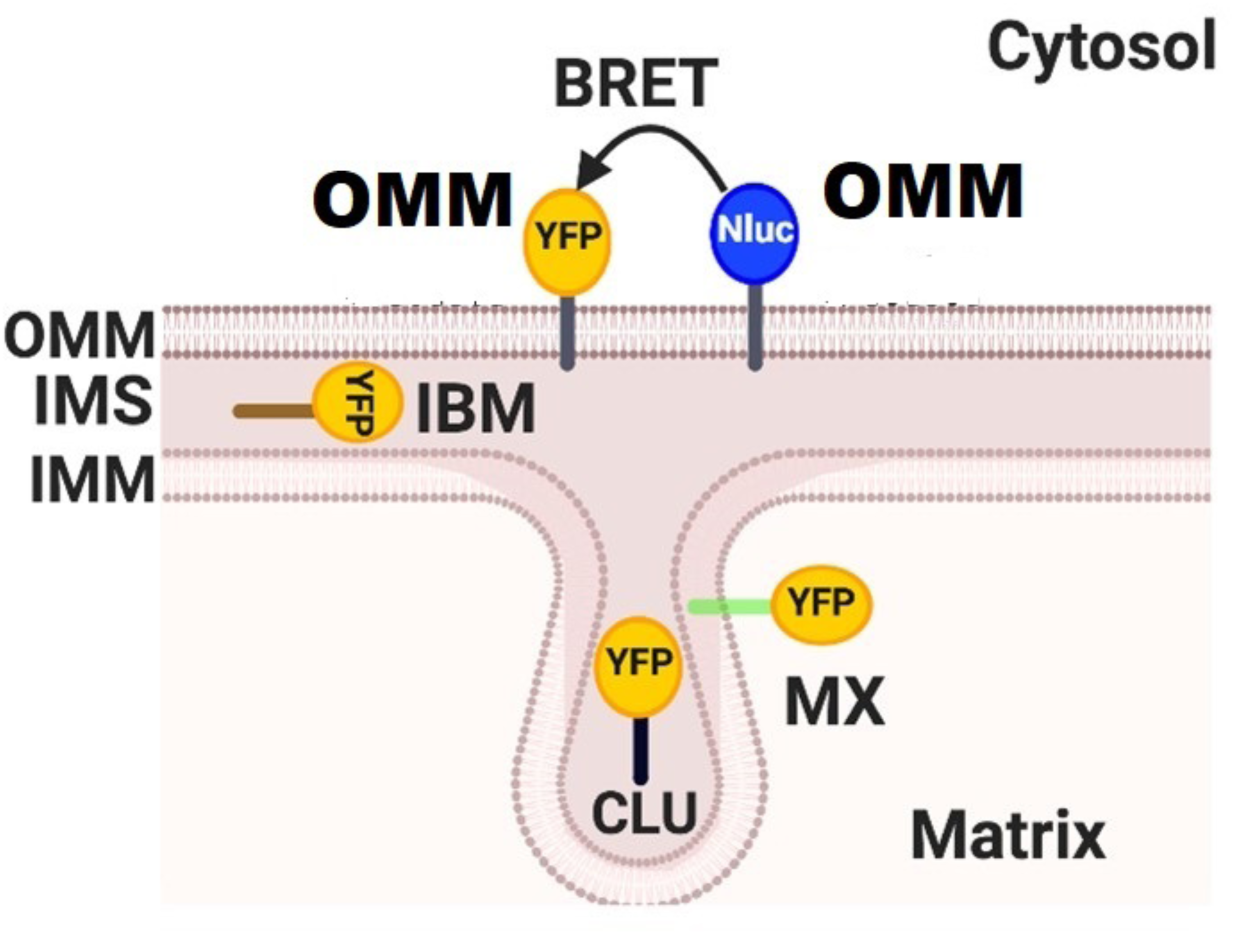
Scheme representing the mitochondrial localization of the different localization sensors. Bioluminescence Resonance Energy Transfer, BRET; Cristae Lumen (CLU); Inner Boundary Membrane (IBM); Inner Mitochondrial Membrane (IMM); Inter-Membrane Space (IMS); matrix (MX); Nanoluciferase, Nluc; Outer Mitochondrial Membrane (OMM); Yellow Fluorescent Protein, YFP.

**Table 1:**
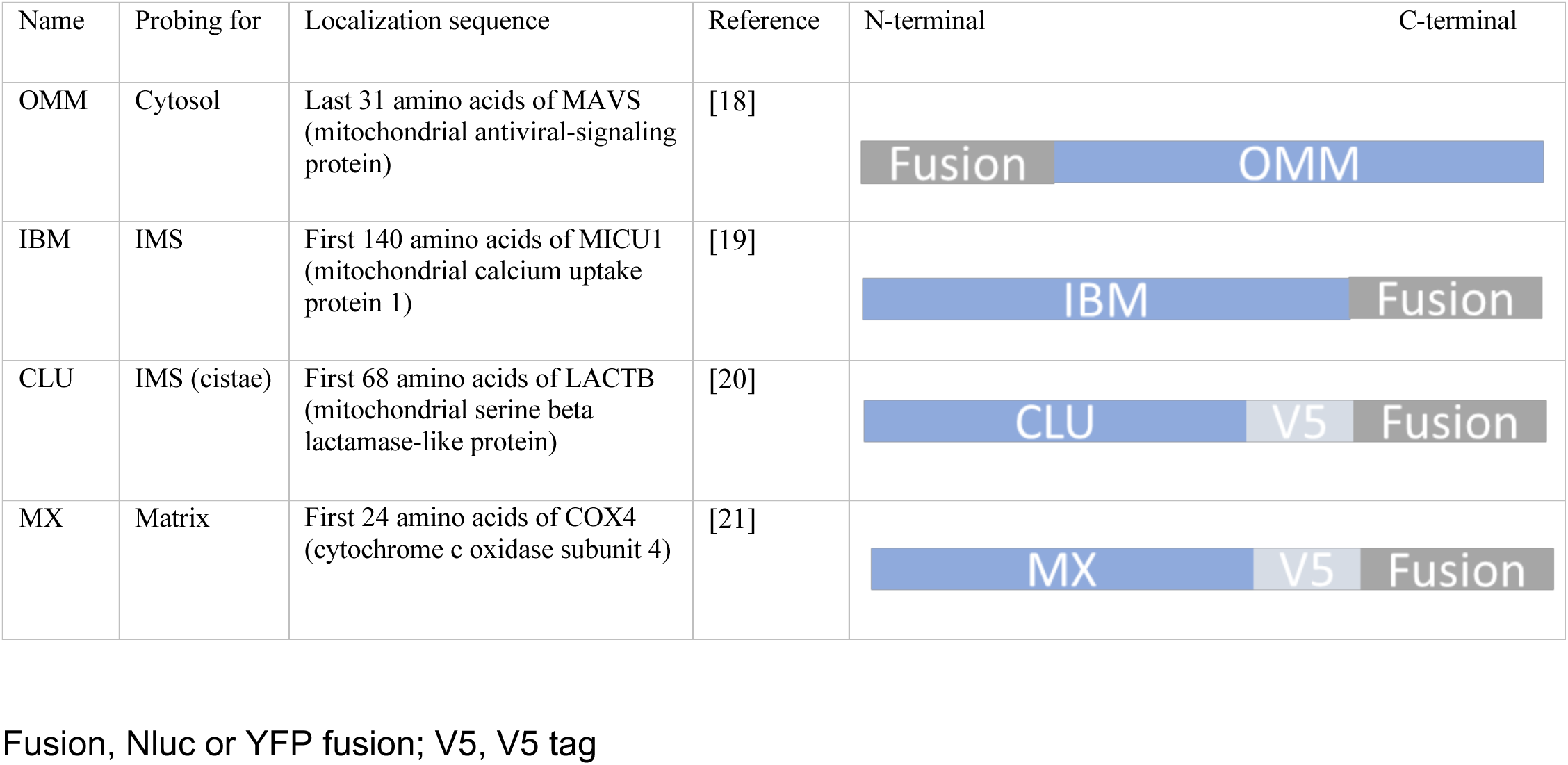
Design of localization sensors.

### Characterization of OMM localization sensor and TOM20 localization

The mitochondrial localization of the YFP-OMM sensor was confirmed by immunofluorescence microscopy and colocalization with the mitochondrial marker protein TOM20 (Figure 2A) in HEK293 cells. Mitochondrial localization of the Nluc-OMM sensor was confirmed by colocalization of the Nluc luminescence signal with Mitotracker fluorescence in live HEK293 cells (Figure 2B). Localization of sensors at specific mitochondrial sub-compartments was studied by BRET by coexpressing different combinations of Nluc and YFP fusion sensors (Figure 2C). Donor and acceptor levels were adjusted for each experimental series to be comparable (Supporting Information Figure S1A,B). Whereas cells coexpressing the OMM sensor fused to Nluc and YFP show a strong BRET signal due to their expected close proximity in the OMM, only marginal signals were observed with the other localization sensors, confirming the specific localization of the Nluc-OMM sensor at the OMM (Figure 2D).

**Figure 2.**
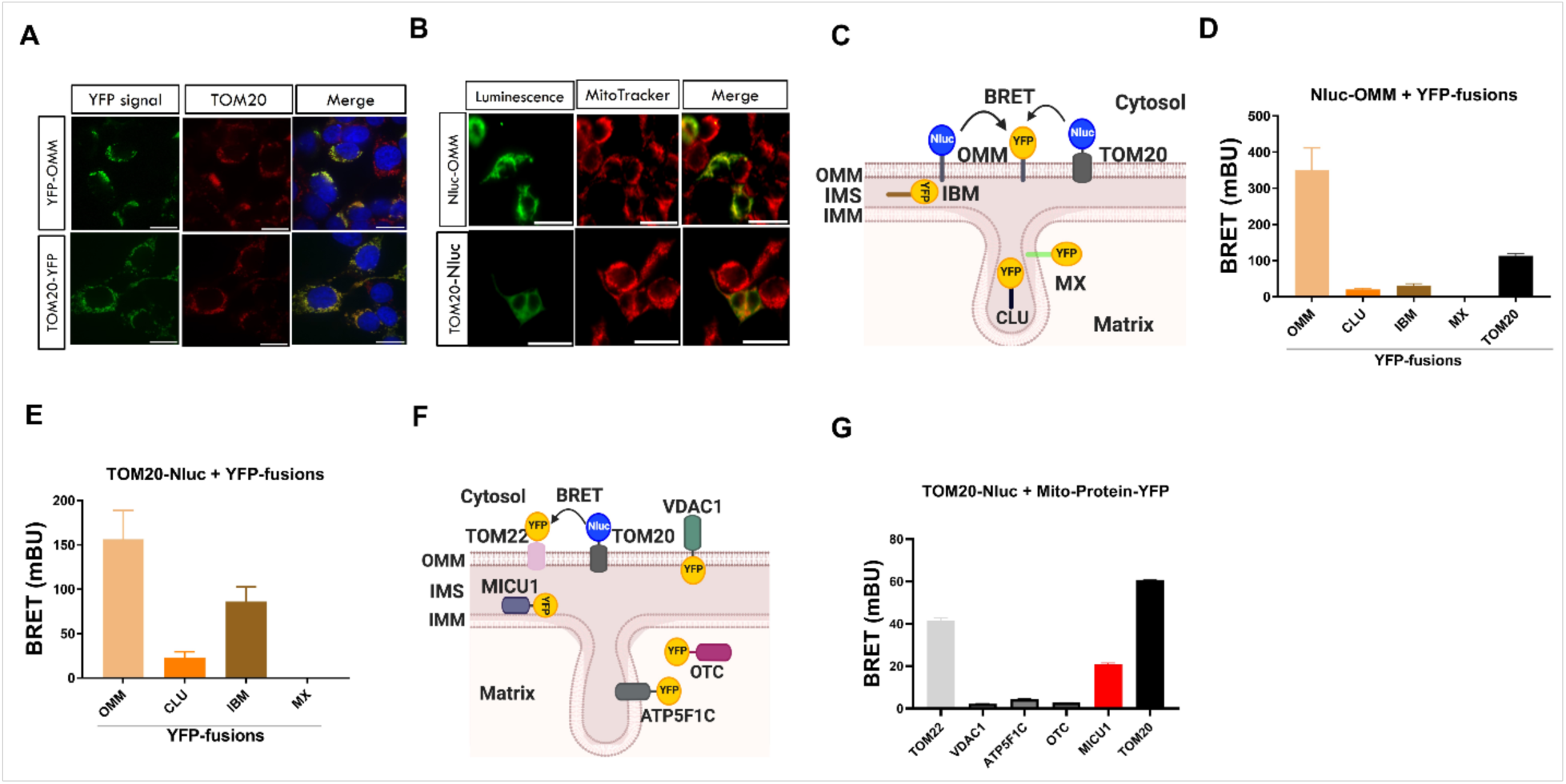
Validation of OMM localization sensor. (A) Immunofluorescence images showing localization of mito-sensors tagged to a YFP (Green) with endogenous TOM20 staining (red) and the superposition of both acquisitions with Dapi (blue). (B) Merged images showing localization of mito-sensors tagged to a Nluc (Green) with Mito Tracker Red CMX Ros staining (Red). (C) Schematic representation of localization sensors in different compartments. (D, E) BRET values represent the proximity between Nluc-OMM, TOM20-Nluc, respectively, and localization sensors (YFP), n =5. (F) Schematic representation of mitochondrial protein sensors in different compartments. (G) BRET values represent the proximity between TOM20-Nluc and mitochondrial proteins (YFP), n =4

We then monitored the molecular proximity between TOM20, a transmembrane protein of the mitochondrial protein translocation machinery, and the OMM sensor. The C-terminal TOM20-YFP fusion protein colocalized as expected with endogenous TOM20 (Figure 2A) and generated a strong BRET signal with the Nluc-OMM sensor (Figure 2D, Supporting Information Figure S1C,D). This result is in agreement with the orientation of the Nluc and YFP moieties towards the cytoplasmic OMM site (Figure 2C). This result was confirmed with the inverse BRET configuration (i.e. TOM20-Nluc/YFP-OMM), excluding any influence of the donor/acceptor fusion (Figure 2B,E, Supporting Information Figure S1C,D). Marginal BRET signals were observed with CLU and MX localization sensors and a significant BRET signal with the IBM sensor (Figure 2E). The latter was unexpected as the YFP moiety of the IBM sensor was designed to probe the IMS and the Nluc moiety of TOM20 to probe the cytosolic side of the OMM (Figure 2C). To better understand the orientation of the TOM20-Nluc fusion protein, we performed coexpression experiments with two additional well-characterized proteins located in the OMM: TOM22, a component of the TOM complex, with the C-terminal YFP fusion predicted to face the cytosol, and the voltage-dependent anion channel 1 (VDAC1) [33] with the C-terminal YFP fusion predicted to face the IMS (Figure 2F,G, Supporting Information Figure S1E,F) [33]. The BRET results are consistent with the orientation of the Nluc fusion of TOM20 towards the cytosol. Marginal BRET signals were observed with two negative control YFP fusion proteins probing the matrix: ATP5F1C (gamma subunit of the mitochondrial ATP synthase) located at the CM, and OTC (ornithine transcarbamylase) located in the matrix. These results exclude thus any other spurious localization of the TOM20-Nluc fusion protein (Figure 2F,G). As the IBM sensor contains the first 140 amino acids of the mitochondrial calcium uptake protein 1 (MICU1), we sought to replicate the IBM result with the full-length MICU1-YFP fusion protein. As for the IBM sensor, a strong BRET signal was observed for the TOM20-Nluc/MICU1-YFP pair, excluding an artefact of the engineered IBM sensor (Figure 2F,G). According to the architecture of the TOM complex, TOM20 forms di(tri)mers [34], a property that was used as positive control to validate the functionality of the TOM20-Nluc fusion protein (Figure 2G). Collectively, these results confirm the expected localization and orientation of TOM20 in the OMM with its C-terminus pointing towards the cytosol. The BRET signal observed with the MICU1 and the derived IBM sensor is most likely due to their proximity during the mitochondrial import process. Indeed, MICU1 is known to be directed towards the IMS via the translocase of the OMM (TOM) complex [35],[36] and to form a stable complex with the TOM complex that can be isolated by co-immunoprecipitation from cardiomyocytes [37]. The absence of significant BRET signals between TOM20 and OTC or ATP5F1C, which are also imported via the TOM complex, is most likely due to a weak and transient interaction, as no such complexes have been captured by co-immunoprecipitation (see Fig. 2G).

### Monitoring the step-by-step translocation of BAX and Drp1 to the OMM

Next, we determined if we could apply the OMM sensor to track the translocation of the pro-apoptotic Bcl-2-associated X Protein (BAX) from the cytosol to the OMM upon apoptosis induction [38,39] (Figure 3A). Based on the detection of caspase 3/7 activity we selected a 4h treatment with 4 µM staurosporine (STS) to induce apoptosis in our HEK293 cells (Supporting Information Figure S2A). Under these conditions, we observed a significant increase in BRET signal between BAX-YFP and the Nluc-OMM localization sensor, indicating the translocation of BAX to the OMM upon apoptosis induction (Figure 3B, Supporting Information Figure S2B,C). This effect was time-dependent, up to 10h of STS treatment (Figure 3C). When using the BAX_A24R mutant, we observed no BRET with OMM (Figure 3A,B). This result is consistent with the reported abolishment of its interaction with TOM22 in a cell-free nanodisc-based system, the initial step of efficient BAX association with OMM [40]. STS treatment had no effect on the BRET signal between OMM and MX localization sensors, excluding any non-specific effect of apoptosis induction on the BRET measurement (Figure 3B). Following this initial interaction of BAX with the OMM, BAX is fully integrated into the OMM, and this step could be detected with the IBM sensor probing the IMS at the level of the inner boundary membrane facing the OMM (Figure 3D, Supporting Information Figure S2D,E). Only background BRET signals were observed with the BAX_A24R mutant and the MX and CLU sensors, confirming the specificity of the signal with the IBM sensor (Figure 3D; Supporting Information Figure S2F,G,H). Collectively, these data indicate that our BRET localization sensors detect for the first time in intact cells the initial, TOM22-dependent, step of OMM association of BAX and confirms the importance of the Ala24 of BAX for this interaction in intact cells. The final step, the full membrane integration of BAX is detected with the IBM sensor as the YFP fusion of BAX is now located in the IMS.

**Figure 3.**
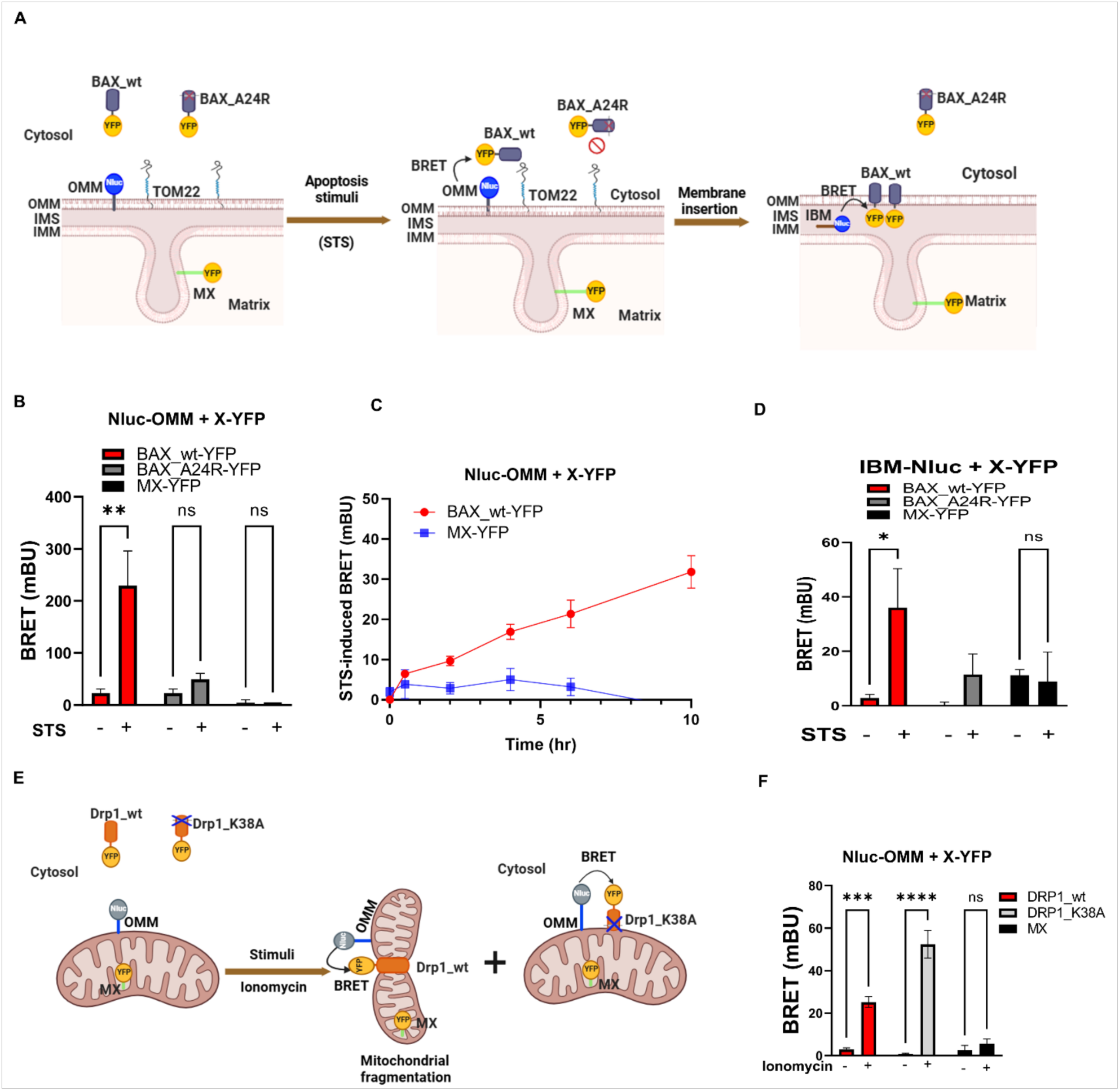
Application of OMM localization sensor. (A) Scheme showing the localization of wild-type BAX (BAX_wt) and the mutant BAX_A24R in the cytosol in healthy and apoptotic HEK293 cells. (B) BRET values representing the proximity between Nluc-OMM and BAX-wt-YFP following apoptosis induced by STS, which was abolished in the presence of BAX_A24R. HEK293 cells were treated or not with STS for 4 hr, followed by BRET measurement n=3. (C) Kinetics of STS-induced BAX_wt translocation from the cytosol to the outer mitochondrial membrane. (D) BRET values representing the proximity between IBM Nluc and BAX-wt-YFP following apoptosis induced by STS, n=5. (E) Scheme showing the localization of wild-type Drp1 (Drp1_wt) and the mutant Drp1_K38A in the cytosol and upon stimulation with ionomycin in HEK293 cells. (F) BRET values representing the proximity between Nluc-OMM and Drp1-wt-YFP and mutant Drp1-K38A-YFP induced by ionomycin treatment, n=3.

We next applied the Nluc-OMM sensor to the detection of the translocation of the Dynamin-related Protein 1 (Drp1), a GTPase regulating mitochondrial fission [41]. Treatment with ionomycin, to raise the intracellular calcium levels, induced a significant BRET signal with wild-type Drp1 (Figure 3E,F, Supporting Information Figure S2I,J). A further increase in BRET signal was observed with the enzymatically dead K38A mutant known to be recruited to mitochondria but not to induce fission (Figure 3F), thus stabilizing the interaction of Drp1 with the OMM [41,42]. Taken together, our sensors reliably monitor the recruitment of cytosolic proteins such as BAX and Drp1 to the OMM in a quantitative manner and confirm the multistep process of OMM integration of BAX as previously suggested by in vitro studies, including the Tom22-dependent step.

### IBM and CLU sensors reliably monitor protein localization in two distinct IMS subcompartments

Next, we validated the mitochondrial localization of the IBM and CLU localization sensors. The respective YFP and Nluc fusion proteins extensively colocalized with mitochondrial markers in HEK293 cells as detected by fluorescence and luminescence microscopy, respectively (Figure 4A,B). To determine their localization in mitochondrial sub-compartments, molecular proximity with different localization sensors was measured by BRET. Strong BRET signals were observed for CLU-Nluc/CLU-YFP and IBM-Nluc/IBM-YFP BRET pairs, as expected, but not for the respective CLU/IBM BRET pairs, nor between IMS sensors and OMM and MX sensors (Figure 4D,E, Supporting Information Figure S3A,B). Collectively, this confirms the exclusive localization of the two IMS sensors in the IMS with distinct and non-overlapping localization of CLU and IBM sensors, most likely corresponding to the crista lumen (CLU) and the IMS between the OMM and the inner boundary membrane (IBM) of the IMM.

**Figure 4.**
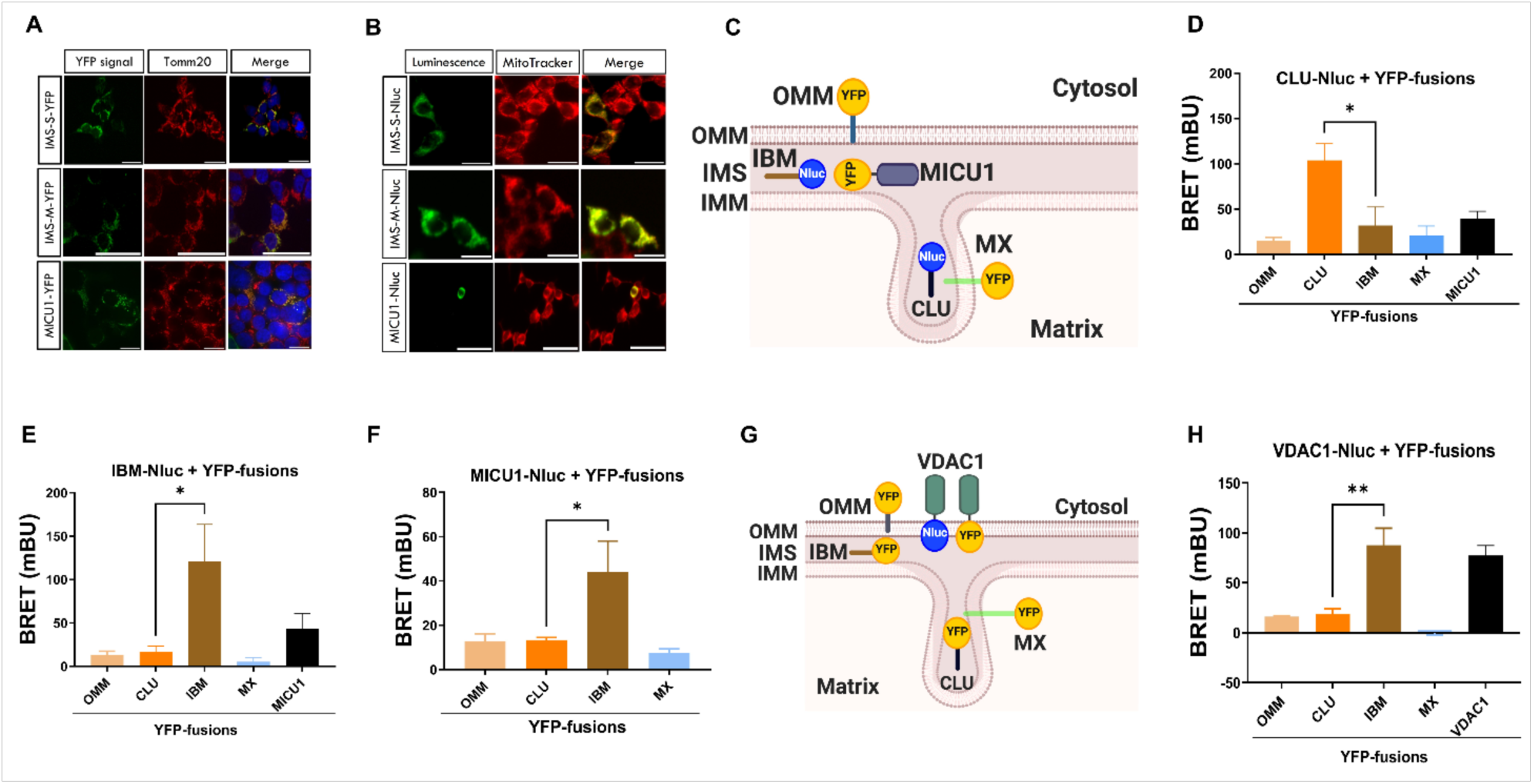
Validation and application of IMS localization sensors. (A) Immunofluorescence images showing mito-biosensors tagged to a YFP (Green) with endogenous TOM20 staining (red) and the superposition of both acquisitions with Dapi (blue). (B) Images showing colocalization of mito-sensors tagged to Nluc (Green) with Mito Tracker Red CMX Ros staining (Red). (C) Schematic representation of mito-biosensors in different compartments. (D,E,F) BRET values represent the proximity between CLU-Nluc, IBM-Nluc, and MICU1-Nluc, respectively, and localization sensors (YFP), n =5. (G) Schematic representation of the localization of VDAC1 and localization biosensors in different compartments. (H) BRET values representing the proximity between VDAC1-Nluc and loc-sensors (YFP), n = 4.

To further explore the differential localization of CLU and IBM sensors, we determined the precise localization of MICU1, a regulatory protein of the MCU (mitochondrial Ca^2+^ uniporter) complex, reported to be exclusively located at the IBM [19]. Mitochondrial localization of MICU1-YFP was first confirmed by fluorescence microscopy (Figure 4A). BRET signals were observed between IBM-Nluc and MICU-YFP and the inverse BRET donor/acceptor configuration (MICU-Nluc/IBM-YFP) but not with the CLU-YFP sensor (Figure 4E,F; Supporting Information Figure S3C,D,E,F). No BRET signal was detected with the OMM and MX sensors (Figure 4E,F), confirming the specificity of the assay. Taken together, the MICU1 results are consistent with its IBM localization and confirm the non-overlapping localization of the CLU and IBM sensors.

Next, we used the IMS sensors to detect the precise mitochondrial localization of VDAC1, described to be located in the OMM with the C-terminus oriented towards the IMS [33]. Consistently, molecular proximity of VDAC1-Nluc was only observed with the IBM YFP-tagged sensors (Figure 4G,H; Supporting Information Figure S3G,H). The functionality of the VDAC1-Nluc fusion protein was further indicated by the detection of previously reported VDAC1 di(oligo)mers [43,44] by BRET (Figure 4G,H). Collectively, these results further validate the distinct localization of the IBM and CLU sensors and show the compatibility of the IBM sensor to measure BRET with OMM proteins with the Nluc/YFP facing the IMS (VDAC1, BAX).

### Characterization and application of the MX localization sensor

The extensive colocalization of the MX sensor with mitochondria labeled with Tom20 antibody or mitotracker confirmed the mitochondrial localization of the MX-YFP and MX-Nluc sensors in fluorescence and luminescence microscopy experiments, respectively (Figure 5A,B). Localization of the MX sensor in mitochondrial subcompartments was probed by BRET with different location sensors (Figure 5C). The expected BRET signal was observed between MX-Nluc and MX-YFP, which was significantly different from BRET signals observed with the other localization sensors, consistent with matrix localization of the MX sensor (Figure 5D, Supporting Information Figure S4A,B).

**Figure 5.**
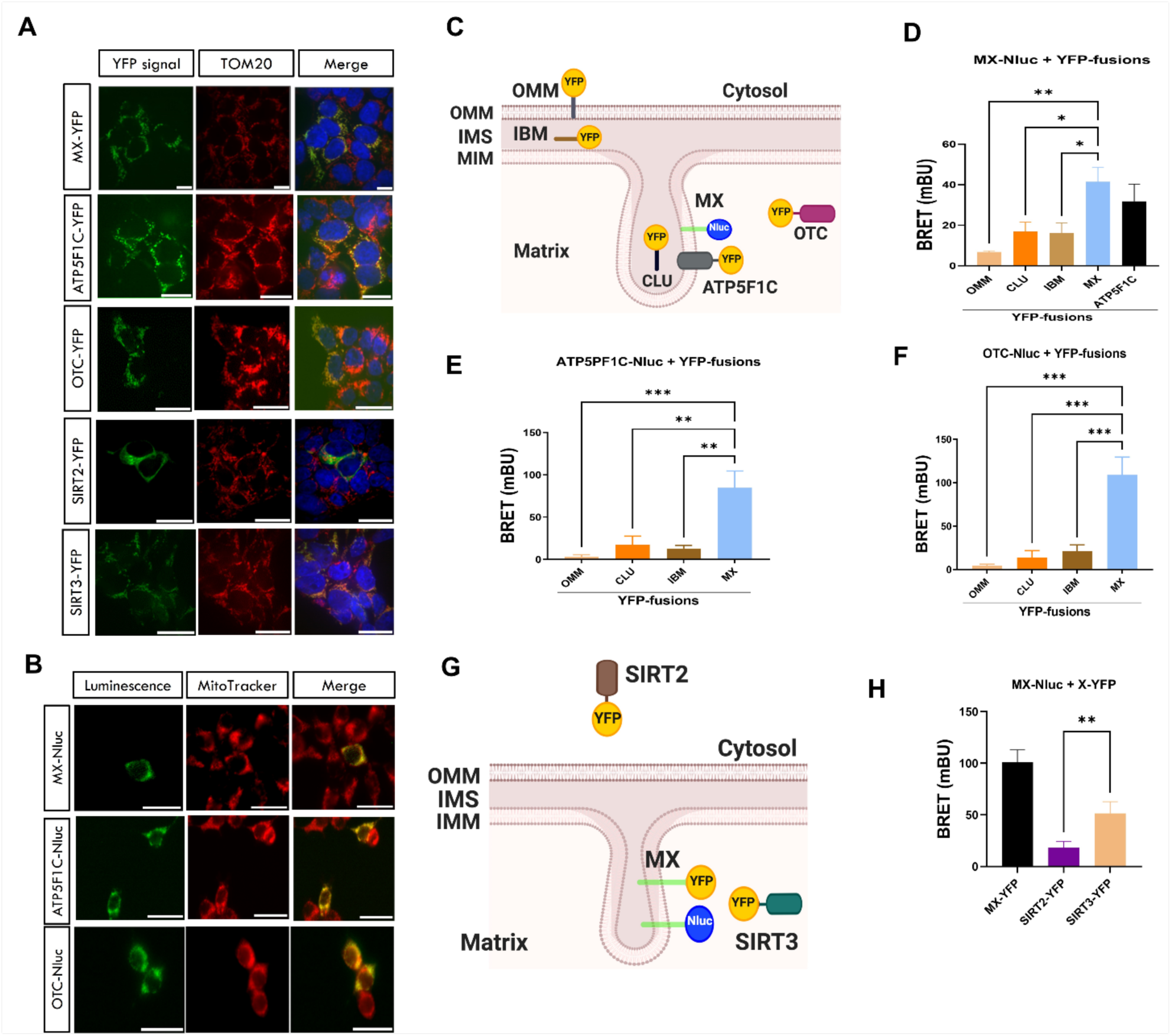
Validation and application of the matrix localization sensor. (A) Immunofluorescence images showing localization of mito-sensors tagged to a YFP (Green) with endogenous TOM20 staining (red) and the superposition of both acquisitions with Dapi (blue). (B) Merged images showing localization of mito-sensors tagged to a Nluc (Green) with Mito Tracker Red CMX Ros staining (Red). (C) Schematic representation of localization sensors in different compartments. (D, E, F) BRET values represent the proximity between -MX-Nluc, ATP5F1C-Nluc, and OTC-Nluc, respectively, and localization sensors (YFP). n =5. (G) Schematic representation of the localization of the Sirtuins and MX loc-biosensor in different compartments. (G) BRET values represent the proximity between MX-Nluc, MX-YFP, and sirtuins (YFP), n = 4

This conclusion was further confirmed by the significant BRET between MX-Nluc and ATP5F1C-YFP located in cristae with the YFP facing the matrix (Figure 4D). In agreement with this result, we also observed a BRET signal in the inverse BRET configuration (ATP5F1C-Nluc/MX-YFP), which was significantly different from the signals observed with all other YFP-tagged localization sensors (Figure 5E; Supporting Information Figure S4C,D). Similar results were obtained with the soluble matrix protein OTC, showing molecular proximity only with the MX sensor (Figure 5F; Supporting Information Figure S4E,F).

Finally, we investigated the ability of the MX sensor to discriminate between the localization of silent information regulator sirtuin (SIRT)2, predominantly located in the cytoplasm [45] and matrix-localized SIRT3 [46] by BRET (Figure 5G). The MX sensor showed BRET signal with SIRT3, which was significantly different from the background signal with SIRT2 (Figure 5H, Supporting Information Figure S4G,H), demonstrating the capacity of the MX sensor to discriminate between the localization of SIRT2 and SIRT3.

## Conclusion

BRET is based on the non-radiative Förster resonance energy transfer that occurs between 1 to 10 nm [22,23]. This range is well-adapted to determine the precise localization of mitochondrial proteins at the level of mitochondrial subcompartments. We developed here a complete set of non-overlapping bystander localization sensors covering all mitochondrial subcompartments, ranging from the OMM to the IMS, IMM and Matrix. Due to the excellent spatial resolution of the sensors, we were also able to distinguish between two structurally distinct IMS subcompartments, the space between the OMM and the IMM, due to a sensor targeted to the IBM and the cristae lumen (CLU) generated by the cristae junction, composed of the mitochondrial contact site and cristae organizing system (MICOS) and optic atrophy 1 (OPA1).

In the case of transmembrane-spanning proteins, BRET may provide additional information on their membrane orientation due to the engineering of the energy donor/acceptor at specific locations of the protein of interest. Results obtained with our BRET localization sensors are consistent with the predicted membrane orientation of TOM20, TOM22, VDAC1, and ATP5F1C, and no alternative membrane orientations were observed for none of them under the current experimental conditions. The unexpected BRET between TOM20 and MICU1 most likely reflects the mitochondrial import of MICU1 through the TOM complex. For MICU1, conflicting localizations either to the IMS or the mitochondrial matrix have been initially reported based on functional studies [47,48], resulting in very different mechanistic models of regulation. Our localization of MICU1 at the IBM facing the IMS is in line with recent APEX electron microscopy (EM) [49] and cryEM studies [50].

Translocation of two cytosolic proteins, BAX and Drp1, to the OMM was detected in intact cells with our BRET localization sensors. In the case of BAX, two different steps could be distinguished: upon STS stimulation, the interaction with the cytosolic face of the OMM, detected with the OMM sensor probing the OMM from the cytosolic site, and the full membrane integration of BAX, detected with the IBM sensor, probing the OMM from the IMS site. Instrumental for capturing these interactions and for defining their specificity was the BAX-A24R mutants reported in previous in vitro studies to be unable to interact with TOM22, the primary mitochondrial target receptor of BAX [40]. Our BRET sensors confirmed these findings now in intact cells and upon stimulation by STS. Of note, only the IBM sensor detected full BAX membrane insertion, but not the CLU sensor probing the second IMS subcompartments at the levels of cristae. These BRET assays have a superior spatial precision, are more quantitative, faster, and more suitable for high-throughput (HTS) applications [22] compared to existing BAX recruitment assays that are based on microscopic or subcellular fractionation approaches [51,52].

The OMM sensor was also able to monitor the recruitment of Drp1 to the OMM. This is remarkable, knowing that Drp1 recruitment promotes mitochondrial fission at the level of the OMM in a highly dynamic manner. Application of the GTPase-defective Drp1-K38A mutant, known to be recruited to the OMM without inducing fission, indeed improved the BRET signal, most likely by protecting the OMM from fission.

Limitations should be considered. An inherent limitation of energy transfer-based techniques, such as BRET, is the use of fusion proteins between the proteins of interest and BRET energy donors and acceptors. Their impact can be minimized by using Nluc, which is the smallest available energy donor with 19 kDa as compared to *Renilla* luciferase (36kDa) and Firefly luciferase (61 kDa) [17]. As the modifying proteins can potentially compromising protein function or inducing mislocalization, fusion proteins should be carefully characterized. Overexpression of proteins is another concern that should be carefully controlled. Use of the highly sensitive Nluc energy donor (approximately 50 times brighter than *Renilla* luciferase) allows to scale down the expression levels of fusion proteins [17]. Tom20 and Tom22 are part of the Tom complex of the protein import machinery through the OMM [34]. Based on our results we do not think that the current sensors are adapted to measure mitochondrial protein import through proximity with Tom20 and Tom22. These interactions are either too transient and/or the molecular proximity or orientation of the Tom20 and Tom22 fusion proteins are not well adapted. This avoids confounding BRET signals due to proximity during protein import thus increasing the specificity of our BRET assays. Whether the interaction of Tom20 and MICU1 is the exception remains to be clarified in a more dedicated study. To measure mitochondrial protein import on a general scale, most likely, more dedicated BRET assays have to be designed.

## Materials and Methods

### Plasmid constructs

OMM localization sensors were generated through the fusion of the last 31 amino acids of the mitochondrial antiviral signaling protein (MAVS) (Addgene # 79056) to the N-terminus of the yellow fluorescent protein (YFP) (YFP-OMM) or NanoLuc (Nluc) (Nluc-OMM). IMS localization sensors are referred to here as IBM and CLU. IBM localization sensors were generated through the fusion of the first 140 amino acids of the mitochondrial calcium uptake protein 1 (MICU1), including the poly-basic domain necessary for IBM localization (Addgene # 79057) in front of YFP (IBM-YFP) or Nluc (IBM-Nluc). CLU localization sensors were generated through the fusion of the first 68 amino acids of the mitochondrial serine beta-lactamase-like protein (LACTB) (Addgene # 79058) in front of YFP (CLU-YFP) or Nluc (CLU-Nluc). MX localization sensors were engineered by fusing the first 24 amino acids of cytochrome c oxidase subunit 4 (COX4) (Addgene # 49151) in front of YFP (MX-YFP) or Nluc (MX-Nluc) (Table 1).

TOM20-YFP(Nluc) fusion proteins were obtained by fusing the TOM20 coding region (Addgene # 55146) in its c-terminus to YFP or Nluc. A similar strategy was applied to TOM22 (Addgene # 111697), ATP5F1C (Addgene # 108933), ornithine transcarbamylase (OTC) (Addgene # 71877), and MICU1 (Addgene # 79057) YFP and Nluc fusion proteins. The Plasmid coding for pEYFP-C1-Drp1 was obtained from Addgene (Addgene # 45160), and the pEYFP-C1-Drp1-K38A mutant was engineered from pEYFP-C1-Drp1 through site-directed mutagenesis using the QuikChange Lightning Multi Site-Directed Mutagenesis Kit (Agilent Technologies, Santa Clara, CA).

The cDNAs encoding human BAX, VDAC1, SIRT2, and SIRT3 with c-terminal GFPSpark® tags were purchased from Sino Biological, Inc. (China). The BAX-A24R mutant was generated through site-directed mutagenesis using the QuikChange Lightning Multi Site-Directed Mutagenesis Kit (Agilent Technologies, Santa Clara, CA). All plasmid constructs were verified by sequencing (Eurofins).

### Cell culture and transfection

The Human embryonic kidney cells (HEK293T) cells were grown in a complete medium (Dulbecco’s modified Eagle’s medium containing 10% (v/v) fetal bovine serum (FBS), 4.5g/L glucose, 100 U/mL penicillin, 0.1 mg/mL streptomycin, and 1mM glutamine) (Invitrogen, CA, USA). For all experiments, cells were maintained at 37 °C with 95%O2, and 5% CO2. For transfection, cells were transiently transfected with JetPEI transfection reagent according to the supplier’s instructions (Polyplus-transfection, New York, NY).

### Immunofluorescence microscopy

HEK293T cells at a density of 1x10^4^ per well were plated in a 24-well cell culture plate on a previously coated poly-L-lysine glass cover slip and maintained overnight at 37°C, 5% CO2. Then, cells were transfected with the YFP fusions of the localization sensors as mentioned above and incubated for a further 24 h. Cells were washed with PBS, fixed in 2% paraformaldehyde for 10 min at room temperature, and washed three times with PBS for 5 min. Cells were permeabilized with 0.1% Triton X-100 for 5 min, washed three times with PBS for 5 min, and incubated with blocking buffer (3% BSA in PBS, 0.1% Tween) for 1 h. Then, cells were washed three times with PBS for 5 min, and incubated with anti-TOM20 antibody (1:1000) (Alexa Fluor 647, Abcam, Cambridge, UK) for 1 h in blocking buffer. To stain the nuclei, cells were incubated with DAPI (Santa Cruz Biotechnology, Dallas, TX, USA) diluted 1:1000 in PBS for 5 min. Then, cells were washed twice with PBS for 5 minutes and the slides were mounted with glycergel mounting medium (Dako, Agilent, USA) and images were acquired under 100×oil objective using Leica DMi6000 inverted Fluorescent microscope (Leica, Germany) with excitation/emission (ex: 491 / em: 525 nm) filters to image the YFP fusion loc-sensors, (Ex: 633nm, Em: 694 nm) for TOM20 and (ex: 405 nm / em: 447 nm) for DAPI. Images were finally analyzed using Fiji [31].

### GloMax Galaxy Bioluminescence Imaging

HEK293T cells at a density of 1x10^4^ per well were plated in a 24-well cell culture plate. Then, cells were transfected with the Nluc fusion proteins of the localization sensors as mentioned above and incubated for a further 24 h. Cells were trypsinized, resuspended in DMEM, and redistributed into 0.1% gelatin-coated µ-slide-8 well high Ibi treat coverslips and further incubated at 37°C, 5% CO2 for 24 h. Then, the media was removed and replaced with a pre-warmed solution of 50 nM MitoTracker™ Red CMXRos (Invitrogen,Waltham, Massachusetts, U.S.) in Opti-MEM and incubated for 12 min at 37°C, 5% CO2. Then, the MitoTracker™ Red CMXRos was removed, and cells were washed once with pre-warmed Opti-MEM. Brightfield, bioluminescence and fluorescence images were acquired sequentially with GloMax Galaxy Bioluminescence Imager (Promega) following the addition of NanoGlo live substrate (Promega) using the following settings: brightfield,100µs and 20% power; fluorescence, excitation: 560/40nm, emission 600LP set at 20% power for 0.5 sec; and total luminescence (without filters) for 2 min. Images were acquired with the auto setting on the GloMax Galaxy left on and finally analyzed using Fiji software [31].

### Apoptosis induction

Transiently transfected HEK293T cells were trypsinized 24 h post-transfection and plated onto poly-L-lysine-coated 96-well plate. Cells were treated with 4µM staurosporine (STS) (MedChemExpress, Sweden) for 4 h before BRET measurements. Apoptosis induction in HEK293T cells was quantified using Caspase-Glo® 3/7 Assay kit (Promega).

### Bioluminescence Resonance Energy Transfer (BRET) Measurement

BRET was performed as previously described [32]. Briefly, 80% confluent HEK293T cells seeded in 24-well plates were transiently transfected with the YFP and/or Nluc-fusion proteins of the localization sensors or full mitochondrial proteins. Then, 24 h post-transfection, cells were transferred into a precoated poly-L-lysine 96-well white plate (Revvity, Waltham, MA) and incubated for an additional 24 h. For BRET measurements, the media was discarded, cells washed with PBS and coelenterazine h added at 5μM final concentration. Then the recording was made using two filter settings, 480 ± 20 nm for the donor and, 540 ± 25 nm for the acceptor (Mithras LB 940 plate reader, Berthold Technologies). Milli BRET units (mBRET) were calculated by dividing the emission signals of acceptor (YFP) by that of Nluc (donor) multiplied by 1000.

### Luminescence and Fluorescence measurement

The expression level of the Nluc/YFP-fusion proteins was measured after each transfection. The relative luciferase activity (luminescence) for the Nluc-fusions was obtained from the BRET measurement by taking the emission values of the donor at 480 nm. The fluorescence of the YFP-fusions, on the other hand, was measured following the transfer of the transfected cells into a precoated poly-L-lysine 96-well black plate (Revvity, Waltham, MA), and incubated for an additional 24 h. Cells were washed once with PBS, and 100 μl of PBS was added to each well, and fluorescence emission was measured at 535 nm, following the excitation of the YFP-fusions at 480 nm with Mithras LB 940 plate reader (Berthold Technologies).

### Statistical Analysis

Data are presented as mean ± standard error of the mean (SEM). Statistical analysis was performed using Graphpad Prism and assessed by two-way ANOVA followed by Šidák multiple comparisons test (Figure 3B,D,F), or by student’s t-test (Figure 4D,E,F,H) and by one-way ANOVA followed by Dunnett’s multiple comparisons test (Figure 5D,E,F,H). Results are considered significant for p-values less than 0.05 (p < 0.05).

## Associated content

### Data availability statement

All data associated with this study are available in the paper and the Supporting Information.

### Author Contributions

AOA, ML and RAZ designed BRET sensors and performed BRET experiments. EC performed luminescence imaging experiments. JD, BM and RJ supervised research. JD and RJ obtained funding. The manuscript was written by AOA and RJ with the contributions of all other authors. All authors have given approval to the final version of the manuscript.

## Acknowledgment

This work was supported by Agence Nationale de la Recherche (ANR-19-CE16-0025-01 « mitoGPCR » to R.J.), the Fondation pour la Recherche Médicale (Equipe FRM EQU202503020060, to R.J.), the “Projet de recherche international” (PRI) Inserm édition 2023 to R,J, the Institut National de la Santé et de la Recherche Médicale (INSERM) and the Centre National de la Recherche Scientifique (CNRS).

## Notes

All authors declare no competing financial interest.

