## Supporting Figures for "BRET-Based Mitochondrial Subcompartment Localization Biosensors"

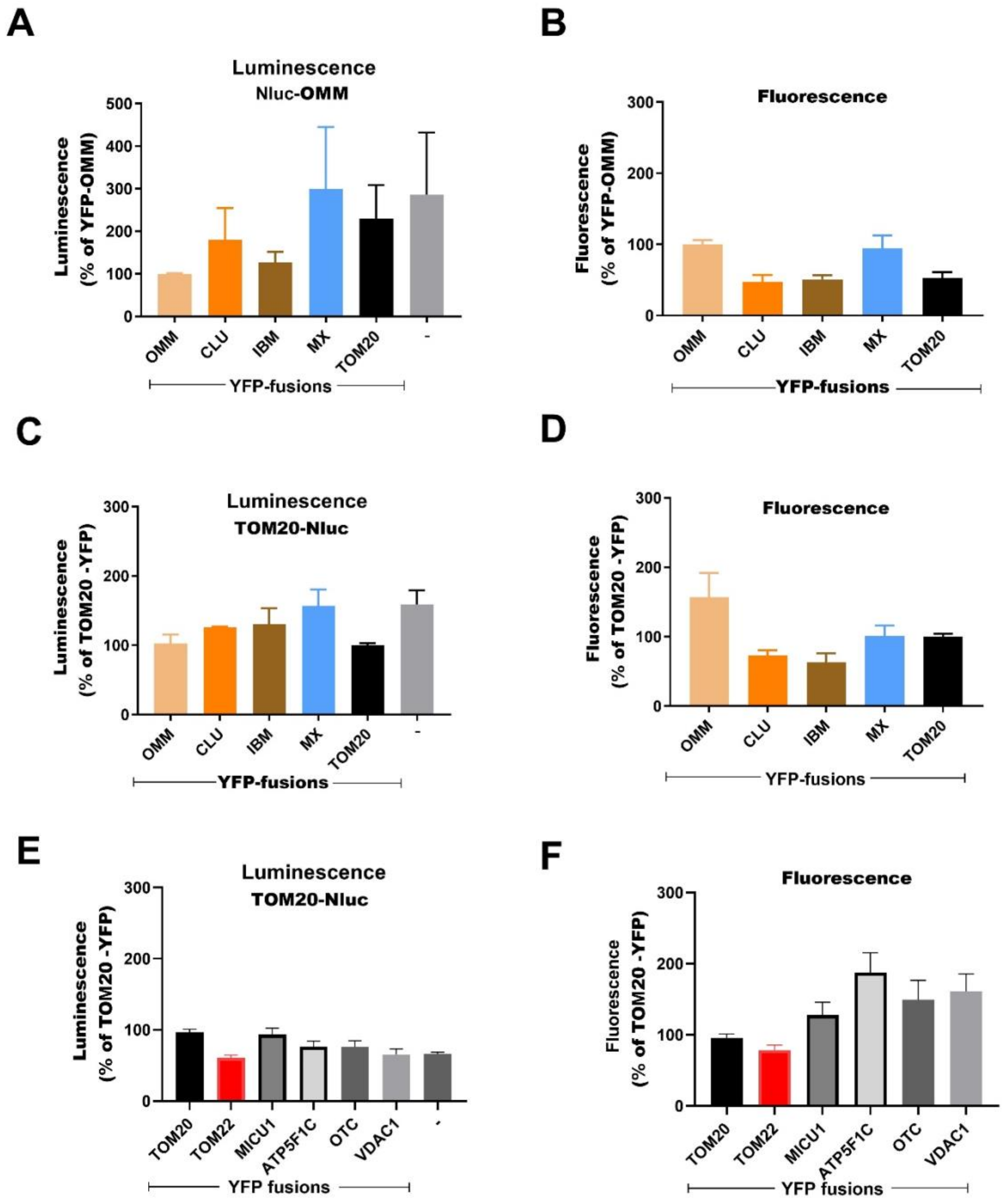


**Figure S1: Relative expression levels of Nluc and YFP fusion proteins.** (A,B) Relative luminescence and fluorescence signals of Nluc-OMM + indicated YFP-fusions. (C,D) Relative luminescence and fluorescence signals of TOM20-Nluc + indicated YFP-fusions. (E,F) Relative luminescence and fluorescence signals of TOM20-Nluc + indicated YFP-fusions. Luminescence values (donor expression levels) were derived from BRET experiments, fluorescence values (acceptor expression levels) were obtained by exciting YFP-fusions directly at 480 nm. Data were normalized to one of the fusion proteins to be able to compare different experiments. Data are mean ± SEM for four to five independent experiments performed in triplicate. See Figure 2 for related BRET values.

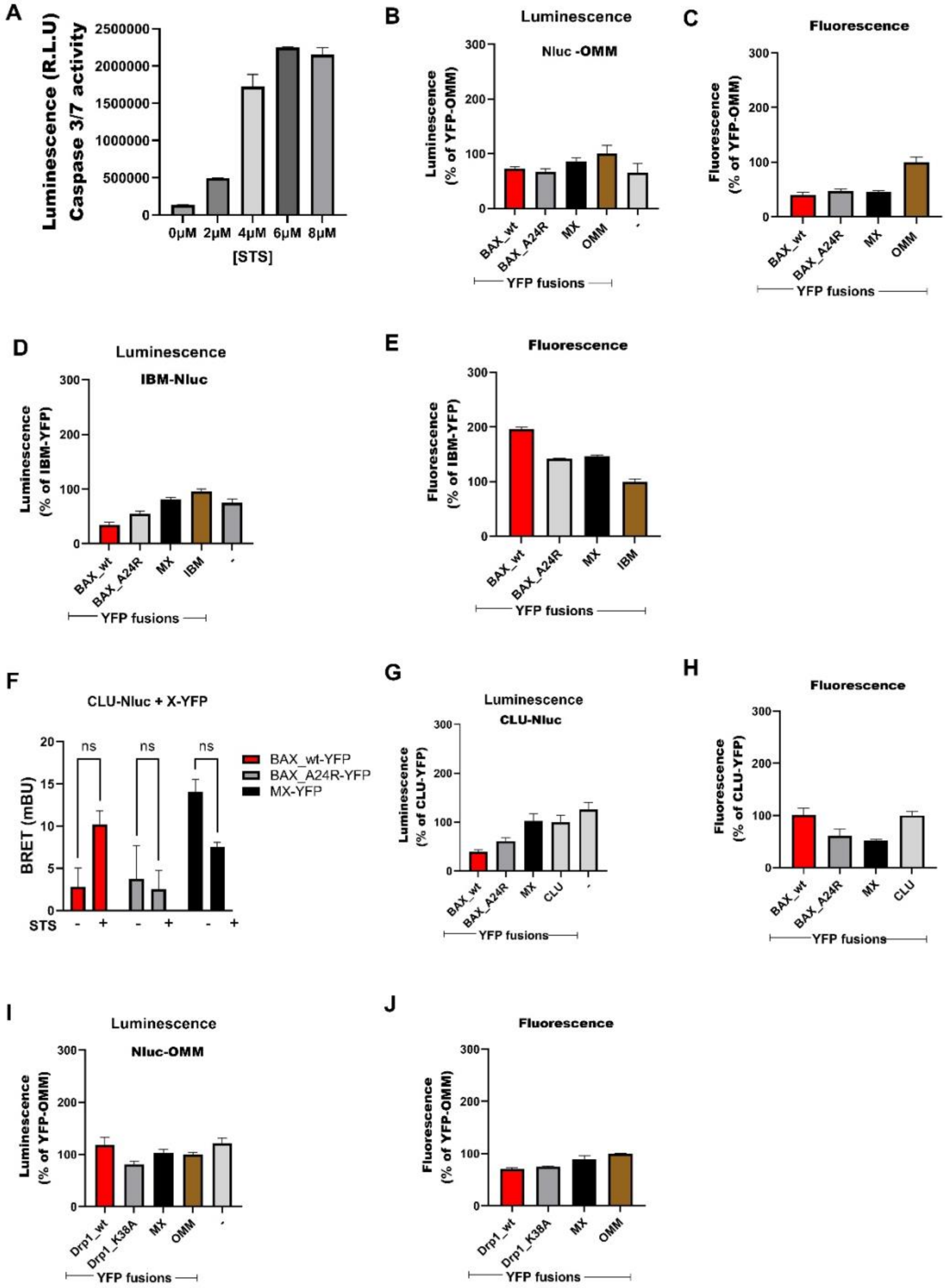


**Figure S2: Apoptosis assay and the relative expression levels of Nluc and YFP fusion proteins.** (A) Apoptosis quantification following the treatment of HEK293T cells with different concentrations of STS for 4 hr. (B,C) Relative luminescence and fluorescence signals of Nluc-OMM and indicated YFP-fusions. (D,E) Relative luminescence and fluorescence signal of IBM-Nluc and indicated YFP-fusions. (F) BRET values representing the proximity between CLU-Nluc and BAX-wt-YFP following apoptosis induced by STS. (G,H) Relative luminescence and fluorescence signal of CLU-Nluc and indicated YFP-fusions. (I,J) Relative luminescence and fluorescence signals of Nluc-OMM and indicated YFP-fusions. Luminescence values (donor expression levels) were derived from BRET experiments, fluorescence values (acceptor expression levels) were obtained by exciting YFP-fusions directly at 480 nm. Data were normalized to one of the fusion proteins to be able to compare different experiments. Data are mean ± SEM for three to five independent experiments performed in triplicate. See Figure 3 for related BRET values.

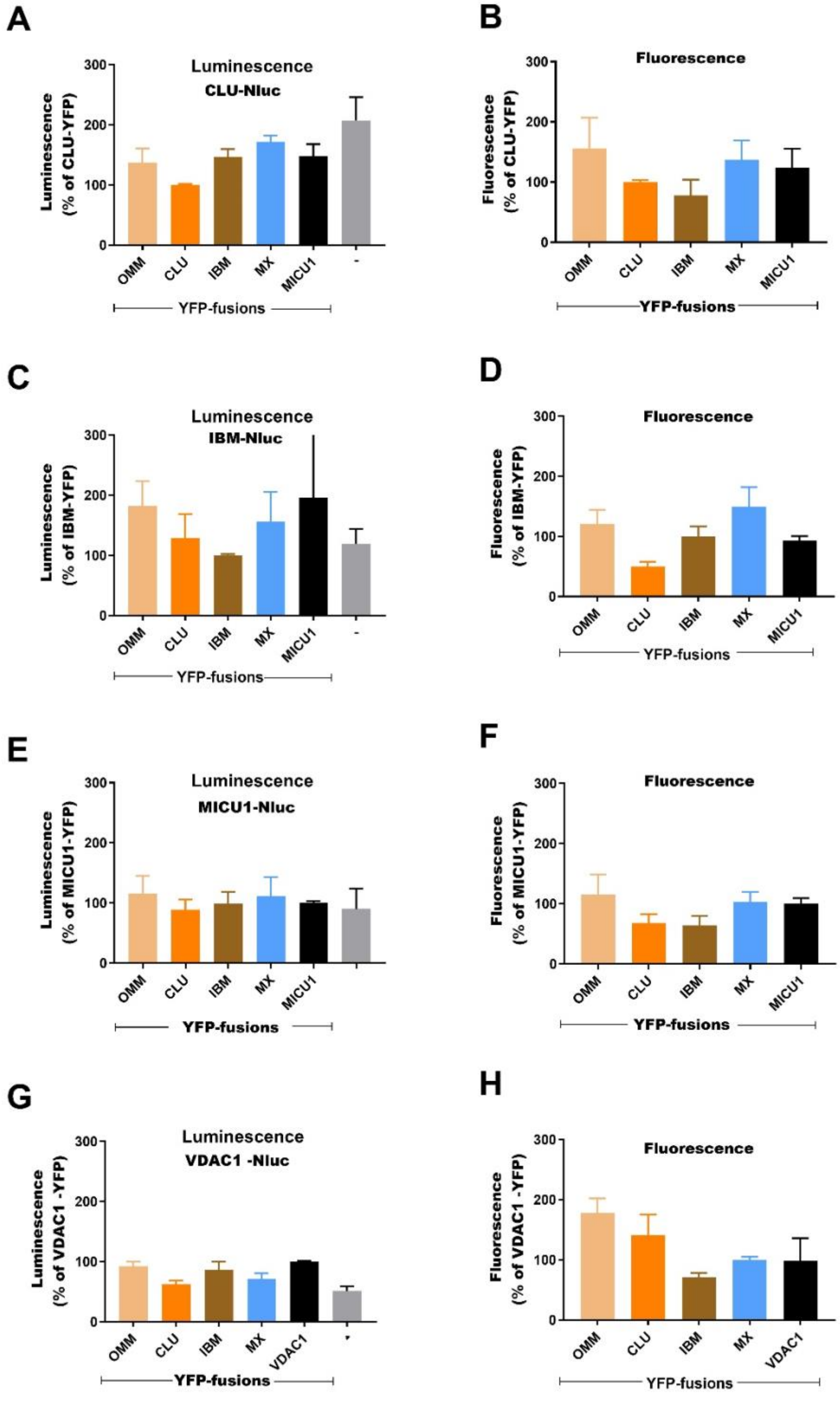


**Figure S3: Relative expression levels of the Nluc and YFP-fusion proteins.** (A,B) Relative luminescence and fluorescence signals of CLU-Nluc and indicated YFP-fusions. (C,D) Relative luminescence and fluorescence signals of IBM-Nluc and indicated YFP-fusions. (E,F) Relative luminescence and fluorescence signals of MICU1-Nluc and indicated YFP-fusions. (G,H) Relative luminescence and fluorescence signals of VDAC1-

Nluc and indicated YFP-fusions. Luminescence values (donor expression levels) were derived from BRET experiments, fluorescence values (acceptor expression levels) were obtained by exciting YFP-fusions directly at 480 nm. Data were normalized to one of the fusion proteins to be able to compare different experiments. Data are mean ± SEM for four to five independent experiments performed in triplicate. See Figure 4 for related BRET values.

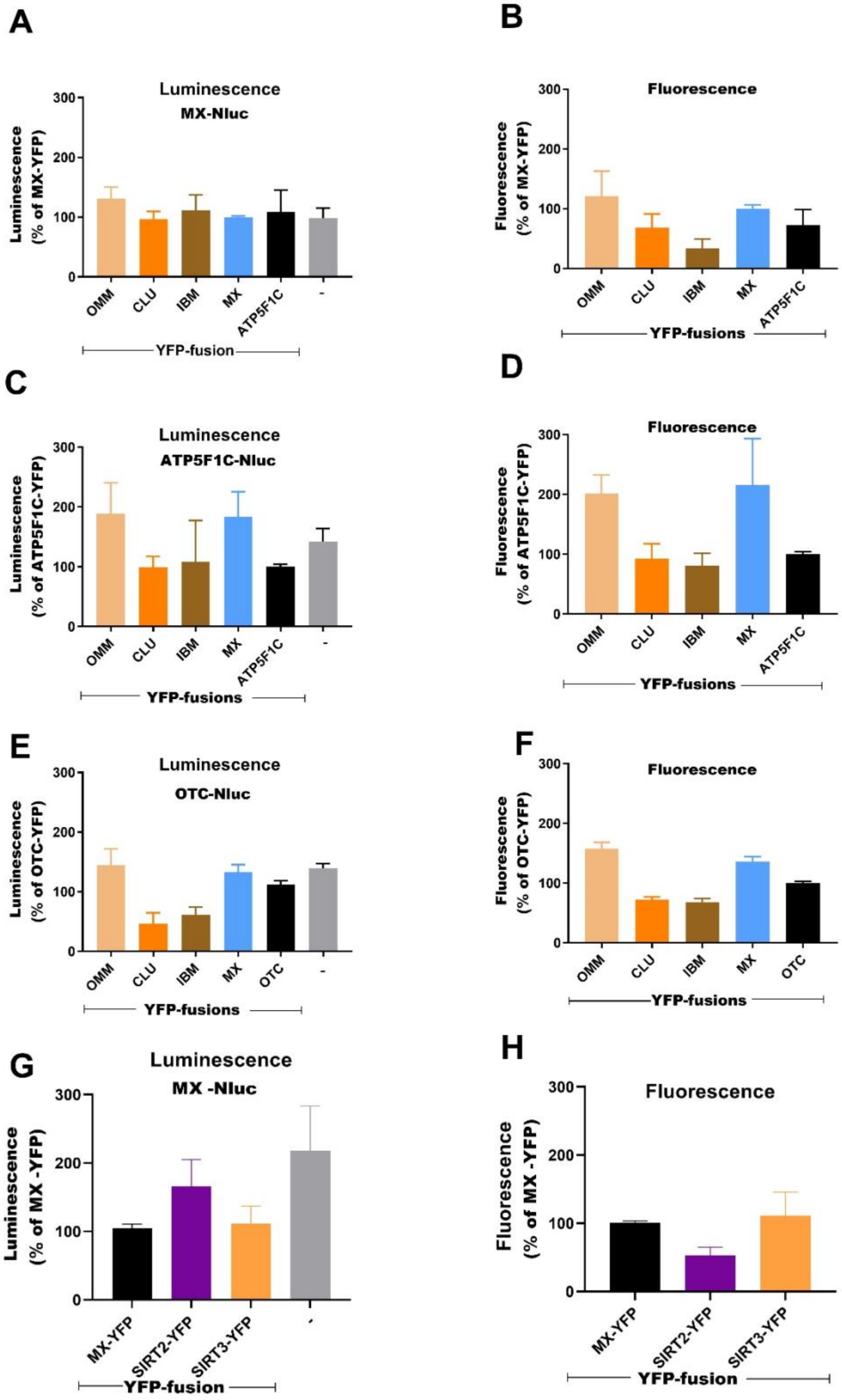
A
Luminescence
MX-Nluc
Luminescence (% of MX-YFP)
0
100
200
300
OMM
CLU
IBM
MX
ATP5F1C
-
YFP-fusion
B
Fluorescence
Fluorescence (% of MX-YFP)
0
100
200
300
OMM
CLU
IBM
MX
ATP5F1C
YFP-fusions
C
Luminescence
ATP5F1C-Nluc
Luminescence (% of ATP5F1C-YFP)
0
100
200
300
OMM
CLU
IBM
MX
ATP5F1C
-
YFP-fusions
D
Fluorescence
Fluorescence (% of ATP5F1C-YFP)
0
100
200
300
OMM
CLU
IBM
MX
ATP5F1C
YFP-fusions
E
Luminescence
OTC-Nluc
Luminescence (% of OTC-YFP)
0
100
200
300
OMM
CLU
IBM
MX
OTC
-
YFP-fusions
F
Fluorescence
Fluorescence (% of OTC-YFP)
0
100
200
300
OMM
CLU
IBM
MX
OTC
YFP-fusions
G
Luminescence
MX -Nluc
Luminescence (% of MX -YFP)
0
100
200
300
MX-YFP
SIRT2-YFP
SIRT3-YFP
-
YFP-fusion
H
Fluorescence
Fluorescence (% of MX -YFP)
0
100
200
300
MX-YFP
SIRT2-YFP
SIRT3-YFP
YFP-fusion

**Figure S4: Relative expression levels of the Nluc and YFP fusion proteins.** (A,B) Relative luminescence and fluorescence signals of MX-Nluc and indicated YFP-fusions. (C,D) Relative luminescence and fluorescence signals of ATP5F1C-Nluc and indicated YFP-fusions. (E,F) Relative luminescence and fluorescence signals of OTC-Nluc and indicated YFP-fusions. (G,H) Relative luminescence and fluorescence signals of MX-Nluc and indicated YFP-fusions. Luminescence values (donor expression levels) were derived from BRET experiments, fluorescence values (acceptor expression levels) were obtained by exciting YFP-fusions directly at 480 nm. Data were normalized to one of the fusion proteins to be able to compare different experiments. Data are mean ± SEM for four to five independent experiments performed in triplicate. See Figure 5 for related BRET values.
